# Face selective patches in marmoset frontal cortex

**DOI:** 10.1101/2020.04.03.023960

**Authors:** David J. Schaeffer, Janahan Selvanayagam, Kevin D. Johnston, Ravi S. Menon, Winrich A. Freiwald, Stefan Everling

## Abstract

Primates have evolved the ability transmit important social information through facial expression. In humans and macaque monkeys, socially relevant face processing is accomplished via a distributed cortical and subcortical functional network that includes specialized patches in anterior cingulate cortex and lateral prefrontal cortex, regions usually associated with high-level cognition. It is unclear whether a similar network exists in New World primates, who diverged ~35 million years from Old World primates and have a less elaborated frontal cortex. The common marmoset (*Callithrix jacchus*) is a small New World primate that is ideally placed to address this question given the complex social repertoire inherent to this species (e.g., observational social learning; imitation; cooperative antiphonal calling). Here, we investigated the existence of a putative high-level face processing network in marmosets by employing ultra-high field (9.4 Tesla) task-based functional MRI (fMRI). We demonstrated that, like Old World primates, marmosets show differential activation in anterior cingulate cortex and lateral prefrontal cortex while they view socially relevant videos of marmoset faces. We corroborate the locations of these frontal regions by demonstrating both functional (via resting-state fMRI) and structural (via cellular-level tracing) connectivity between these regions and temporal lobe face patches. Given the evolutionary separation between macaques and marmosets, our results suggest this frontal network specialized for social face processing predates the separation between Platyrrhini and Catarrhine. These results give further credence to the marmoset as a viable preclinical modelling species for studying human social disorders.

## Introduction

The circuitry responsible for face processing has been well documented in humans and macaques (Puce et al., 1996; Moeller, 2008; Tsao et al., 2008a; Pitcher et al., 2011; Weiner and Grill-Spector, 2015). Old World primate species seem to share a common architecture of this circuitry, with multiple face-selective patches along the occipitotemporal axis that are functionally connected with a larger face processing network that includes several subcortical areas (e.g., hippocampus, amygdala; Schwiedrzik et al., 2015) and face-selective patches in frontal cortex (Tsao et al., 2008a, 2008b). The face-selective patches in frontal cortex have been implicated in processing social context and orofacial movements in macaques – anterior cingulate cortex and lateral prefrontal cortex are differentially activated when in direct visual contact with the face of a conspecific (Yoshida et al., 2011; Shepherd and Freiwald, 2018). It is unclear whether a similar network exists in New World primates, who separated ~35 million years ago from Old World primates (Schrago and Russo, 2003) and have a less elaborated frontal cortex (Solomon and Rosa, 2014). The common marmoset (*Callithrix jacchus*) is a small New World primate that is ideally placed to address this question given the rich social repertoire inherent to this species (e.g., observational social learning; imitation; cooperative antiphonal calling; Miller et al., 2016). Here, we used ultra-high field (9.4 Tesla) task-based functional magnetic resonance imaging (fMRI) in marmosets to investigate whole brain face processing in marmosets.

Recent fMRI studies have demonstrated that marmosets do indeed possess face patches in a ventral pathway along the temporal lobes that have a similar organization to Old World primates (Hung et al., 2015a, 2015b). Given that marmosets use eye contact and facial expression as a means of social communication (Kemp and Kaplan, 2013; Mitchell et al., 2014; Kotani et al., 2017), we posited that marmosets could also possess a face processing network that extends into frontal cortex. The frontal constituents of face-to-face interaction in macaques have been demonstrated by showing conspecific videos during fMRI acquisition (Shepherd and Freiwald, 2018) – when viewing videos of other macaques in a “direct-gaze” context (i.e., simulated eye contact) a patch of anterior cingulate cortex is differentially activated. Interestingly, this patch is less active when viewing videos of other macaques in an “averted-gaze” context (i.e., while the monkey in the video is looking away). Here, our goal was to employ a marmoset conspecific version of this task during whole brain fMRI to test for the existence of functional face patches in frontal cortex.

By leveraging our recent hardware advances in ultra-stable awake marmoset imaging (Schaeffer et al., 2019a) we acquired whole brain fMRI in four marmosets while they viewed videos of marmoset faces in social (with directed or averted gaze) or non-social (scrambled videos) conditions. To quantify gaze differences between the stimuli, we also performed eye tracking in five marmosets while they performed the same task outside of the MRI environment. To corroborate the connectivity of the task-based circuitry, we utilized our extensive fully awake resting-state fMRI (RS-fMRI) dataset to index functional connectivity. To index structural connectivity, we overlaid the results of tracer-based cellular connectivity data of multiple anterior cingulate cortex injection sites in marmosets (Majka et al., 2020).

## Methods

### Subjects

#### Marmosets

Data were collected from 10 adult marmosets (*Callithrix jacchus;* three female; weight 245-380 g; age 29 - 74 months), with n = 4 for task-based fMRI, n = 5 for RS-fMRI, and n = 5 for the eye tracking outside of the MRI (all but 1 exclusive to the monkeys used in fMRI experiments). Experimental procedures were in accordance with the Canadian Council of Animal Care policy and a protocol approved by the Animal Care Committee of the University of Western Ontario Council on Animal Care. All animal experiments complied with the ARRIVE guidelines.

### Marmoset surgical implantation and head-fixation training

All 10 marmosets underwent an aseptic surgical procedure to implant a head chamber, 5 of which were MRI-compatible (i.e., implanted using non-radio opaque dental cement). The purpose of the chamber was to fix the head and thereby prevent animal motion during MRI acquisition. The chamber implantation procedure is described in detail in Johnston et al. (2018) and Schaeffer et al (2019c); in brief, several coats of adhesive resin (All-bond Universal, Bisco, Schaumburg, Illinois, USA) were applied using a microbrush, air dried, and cured with an ultraviolet dental curing light (King Dental). Then, a two-component dental cement (C & B Cement, Bisco, Schaumburg, Illinois, USA) was applied to the skull and to the bottom of the chamber, which was then lowered onto the skull via a stereotactic manipulator to ensure correct location and orientation. The chamber was 3D printed at 0.25 mm resolution using stereolithography and a clear photopolymer resin (Clear Resin V4; Form 2, Formlabs, Somerville, Massachusetts, USA).

Before MRI acquisition, the marmosets were first acclimatized to the head-fixation system and a mock MRI environment (including sequence sounds; see Schaeffer et al., 2019c for open-source hardware designs for the animal holder and detailed training procedures). Each marmoset was acclimatized over the course of three weeks prior to imaging.

### Face processing task

A block design was used in which nine baseline blocks (18 sec each) were alternated with eight task blocks (12 sec each; see Figure 1). During baseline blocks, a 0.36° circular black cue was displayed in the center of the screen against a gray background – this was in part to be used as a fixation point (although marmosets did not fixate *per se*), but also as a visual stimulus that served to mitigate visual nystagmus invoked by the high magnetic field. During task blocks, the dot disappeared and a video was presented in the center of the screen (6° height × 10.6° width) – 3 stimulus sets were used (counterbalanced between animals), with four pseudo-randomized task conditions each (directed gaze, averted gaze, and scrambled versions of each; see Figure 1 for representative stimuli). For the directed and averted gaze videos, 5 marmosets were filmed while they sat non head-fixed in a marmoset chair (Johnston et al., 2018); 12 sec clips were created using custom video-editing software (iMovie, Apple Incorporated, California, USA). Scrambled versions of the videos were created by random rotation of the phase information using a custom program (Matlab, The Mathworks, Matick, MA) – the same random rotation matrix was used for each frame to preserve motion components.

**Figure 1.**
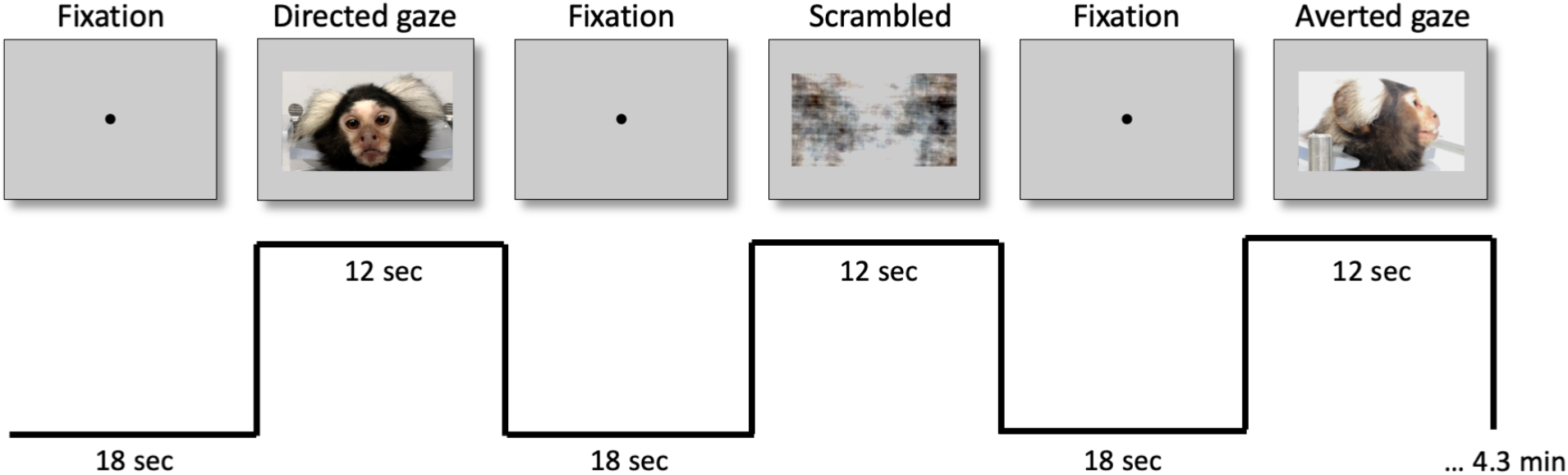
Stimuli and experimental design for task-based fMRI. Top shows the stimuli for the fixation condition and the task conditions, including directed gaze, averted gaze, and scrambled videos. The black line below the stimuli shows the task timing.

Two of the four monkeys were rewarded at the start and end of every block to keep these animals awake and engaged. The liquid reward (diluted sweetened condensed milk) was delivered via infusion pump (Model NE-510, New Era Pump Systems, Inc., Farmingdale, New York, USA). The reward volume was set to 50 microliters per dispense and was delivered over the course of 1 second; note that reward tube was placed outside of the marmosets mouth (~ 5 mm away) and thus they needed to extend their tongue in order to lick the reward from the tube. We have previously isolated the reward-related circuitry using a similar block design paradigm (Schaeffer et al., 2019b; Figure 6 therein) and this circuitry is, for the most part, discrete from the face circuity of interest, as described below.

The stimuli were presented via projector (Model VLP-FE40, Sony Corporation, Tokyo, Japan), reflected from a first surface mirror, which back-projected the image onto a plastic screen that was affixed to the front of the scanner bore. The stimuli were presented via Keynote (Version 7.1.3, Apple Incorporated, California, USA), with the stimulus timing (based on a per image volume repetition time (TR) transistor-transistor logic (TTL) pulse) achieved using a Raspberry Pi (Model 3 B+, Raspberry Pi Foundation, Cambridge, UK) programmed in-house.

### Image acquisition

#### Animal holder and MRI hardware

An integrated animal holder and 5-channel radiofrequency receive array was used to rigidly fix the animal’s head chamber to the receive coil. This hardware was designed in-house and is described in detailed in Schaeffer et al. (2019c), wherein the open-source computer-aided design files are linked. An MRI-compatible camera (Model 12M-i, MRC Systems GmbH, Heidelberg, Germany) allowed for continuous monitoring by a veterinary technician for any sign of struggle or discomfort. Given that skull-attached chambers are generally accompanied by magnetic-susceptibility image artifacts (via differences in the magnetic susceptibility between the chamber, adhesive, air, and tissue, as well as the surgical displacement of the skin, fat, and muscle), we sought to ameliorate this distortion by filling the chamber with a water-based lubricant gel (MUKO SM321N, Canadian Custom Packaging Company, Toronto, Ontario, Canada) prior to each imaging session.

Data were acquired using a 9.4 T 31 cm horizontal bore magnet (Varian/Agilent, Yarnton, UK) and Bruker BioSpec Avance III console with the software package Paravision-6 (Bruker BioSpin Corp, Billerica, MA), a custom-built high-performance 15-cm-diameter gradient coil with 400-mT/m maximum gradient strength (xMR, London, CAN; Peterson et al., 2018), and the receive coil described above. Radiofrequency transmission was accomplished with a quadrature birdcage coil (12-cm inner diameter) built in-house.

#### Imaging parameters

Functional imaging was performed over multiple sessions (days) for each animal, with 6 – 8 task-based functional runs (at 172 volumes each) per animal with the following parameters: TR = 1500 ms, TE = 15 ms, flip angle = 40 degrees, field of view = 64 × 64 mm, matrix size = 128 × 128, voxel size = 0.5 × 0.5 × 0.5 mm, slices = 42, bandwidth = 500 kHz, GRAPPA acceleration factor: 2 (anterior-posterior). T2-weighted structural scans were acquired for each animal during one of the awake sessions with the following parameters: TR = 5500 ms, TE = 53 ms, field of view = 51.2 × 51.2 mm, matrix size = 384 × 384, voxel size = 0.133 × 0.133 × 0.5 mm, slices = 42, bandwidth = 50 kHz, GRAPPA acceleration factor: 2.

### Image preprocessing

The fMRI data was preprocessed using AFNI (Cox, 1996) and FSL (Smith et al., 2004). Raw functional images were converted to NifTI format using dcm2niix (Li et al., 2016) and reoriented from the sphinx position using FSL. The images were then despiked (AFNI’s 3dDespike) and volume registered to the middle volume of each time series (AFNI’s 3dvolreg). The motion parameters from volume registration were stored for later use with nuisance regression. Images were smoothed by a 1.5 mm full-width at half-maximum Gaussian kernel to reduce noise (AFNI’s 3dmerge). An average functional image was then calculated for each session and registered (FSL’s FLIRT) to each animal’s T2-weighted image – the 4D time series data was carried over using this transformation matrix. Anatomical images were manually skull-stripped and this mask was applied to the functional images in anatomical space. The T2-weighted images were then non-linearly registered to the NIH marmoset brain atlas (Liu et al., 2018) using Advanced Normalization Tools (ANTs; Avants et al., 2011) and the resultant transformation matrices stored for later transformation (see below). The olfactory bulb was manually removed from the T2-weighted images of each animal prior to registration, as it was not included in the template image.

### Task-based fMRI comparisons

The task timing was convolved to the hemodynamic response (using AFNI’s ‘BLOCK’ convolution) and a regressor was generated for each condition to be used in a regression analysis (AFNI’s 3dDeconvolve) for each run. All four conditions were entered into the same model, along with polynomial (N = 5) detrending regressors, bandpass regressors, and the motion parameters derived from the volume registration described above. The resultant regression coefficient maps were then registered to template space using the transformation matrices described above and then converted to Z value maps. The Z value maps for each monkey were then compared at the group level via t-test (AFNI’s 3dttest++). To protect against false positives, a clustering method derived from Monte Carlo simulations was applied to the resultant t-test maps (using AFNI’s AlphaSim).

### Resting-state seed analysis

To corroborate the locations of the face patches identified with the task-based analysis, we indexed task-independent functional connectivity of each of the identified temporal face patches described below (i.e., those found by contrasting the directed and averted gaze conditions from the task-based analysis). To do so, we acquire RS-fMRI from five fully awake adult marmosets (including the four marmosets used above) using the same fMRI acquisition parameters as described above, but with 600 volumes. In total, 35 15-minute sessions were acquired and used in this analysis. With this data, we calculated seed-based connectivity across the brain using a 1.5 mm cubic region of interest placed at the center of mass of each face patch in temporal lobes. The preprocessing steps described above were also used for the RS-fMRI seed analysis, apart from the task regressors. Instead, the mean time courses extracted from the four regions of interest were used as the regressors, for each run.

### Comparison with tracer-based cellular connectivity

With the recent release of tracer-based cellular connectivity maps across marmoset cortex in volume space (Majka et al., 2020), we were able to directly compare retrograde histochemical tracing in marmosets with our task-based face selective topologies. Explicitly, focused on the tracer maps from two injections (CJ164-DY and CJ164-FB; marmosetbrain.org for notes on these injections) located most proximally to the cingulate cortex patch found by contrasting the directed and averted gaze task-based fMRI conditions here (i.e., the cingulate patch sensitive to socially-relevant processing). Note that at the time of this experiment (2020), the temporal injections reported by this resource were relatively sparse and not specific to the face patches found with our fMRI design. To bring the fMRI data and tracer-based connectivity into the same space, the anatomical template (described in Majka et al., 2020) was nonlinearly registered to the NIH marmoset brain atlas (Liu et al., 2018) using ANTs (Avants et al., 2011) – the volumetric injection data was then brought into template space via this transformation matrix. Note that the tracer data are for the left hemisphere only; as such, we mirrored the left hemisphere data onto both hemispheres.

### Eye tracking

To investigate differences in patterns of eye movements between conditions we performed eye tracking outside of the scanner (i.e., free of MRI-induced noise, but with identical stimuli; see Figure 1). Eye positions were digitally recorded at 1 kHz via video tracking of the left pupil (EyeLink 1000, SR Research, Ottawa, ON, Canada). Animals were head restrained in a custom chair (Johnston et al., 2018) mounted to a table in a sound attenuating chamber (Crist Instruments Co., Hagerstown, MD, USA). A spout was placed at the monkey’s mouth to deliver reward (acacia gum) via an infusion pump (Model NE-510, New Era Pump Systems, Inc., Farmingdale, New York, USA). In each session, eye position was calibrated by rewarding 300 to 600 ms fixations on dots presented at one of five locations on the display monitor using the CORTEX real-time operating system (NIMH, Bethesda, MD, USA). All stimuli were presented on a CRT monitor (ViewSonic Optiquest Q115, 76 Hz non-interlaced, 1600 × 1280 resolution). A TTL pulse triggered from a photodiode was used to determine the start of each block. Animals were intermittently rewarded at random time intervals to maintain their interest.

Analysis was performed using Python code written in-house. Eye velocity (visual deg/s) was obtained by smoothing and numerical differentiation. Saccades were defined as radial eye velocity exceeding 30 deg/s. Fixations were defined as periods where radial eye velocity remained below 10 deg/s for at least 50 ms. Differences in saccade amplitudes and fixations durations between conditions were analysed with repeated measures analysis of variance (ANOVA) using SPSS (v.25, IBM Corp, 2019) statistical software (post-hoc comparisons were Bonferroni-corrected).

## Results

### Task-based fMRI comparisons

Figure 2 shows group maps comparing the social video conditions (including both directed and averted gaze) and non-social scrambled versions of those videos. As shown in Figure 2, videos containing conspecific faces elicited a broad network that was similar, but stronger than the topology that was elicited using the scrambled versions of the videos. The social conditions showed stronger activation along the occipitotemporal axis with peaks in V4/TEO and TE3. In frontal cortex, the social videos showed peaks laterally in 45/47L and orbitofrontally in 13L. Along the medial cortical surface, the largest differences were present in visual cortex (V1 and V2). We also found subcortical differences between the social and non-social scrambled conditions, including in the superior colliculus (SC), hippocampus (Hipp), Pulvinar (Pul), medial-dorsal nucleus of thalamus (MD), and in the amygdala (Amy).

**Figure 2.**
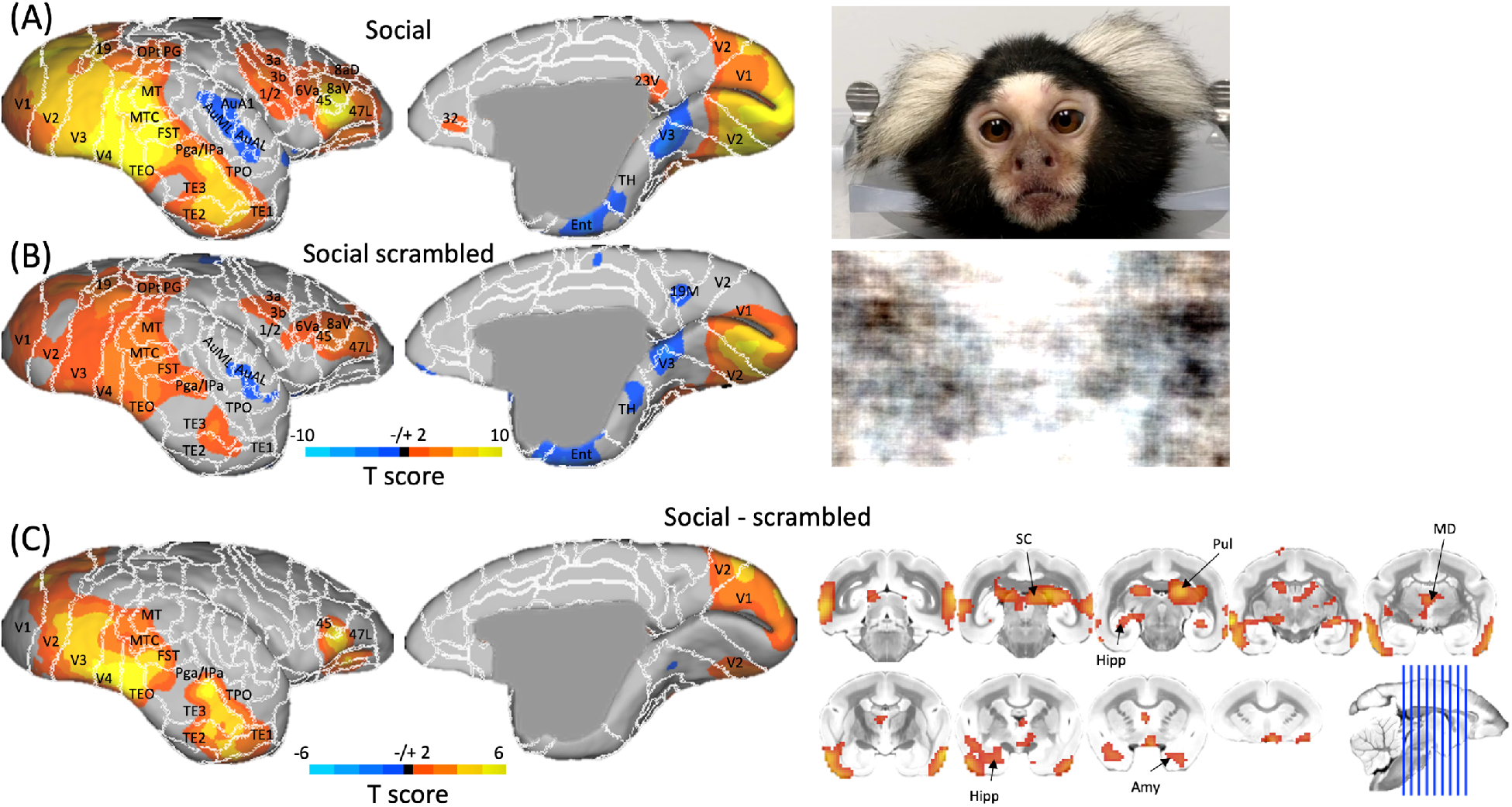
Group functional topology comparisons between social (i.e., directed and averted gaze) and non-social scrambled video conditions displayed on a marmoset cortical surface. (A) shows group topologies for social videos and (B) shows topologies for social scrambled videos, with stills of representative stimuli to the right of each surface. (C) shows the contrast between the social and scrambled conditions, with volumetric display (to show subcortical activation) to the right of the medial and lateral surfaces.

Figure 3 shows the comparison between the two different social conditions – directed gaze and averted gaze. Overall, the broad topologies of these circuitries were similar, but Figure 3C shows several critical differences when contrasting the two conditions. The activation was greater for the directed gaze videos along the occipitotemporal axis – these regions are remarkably similar to the face patches identified in marmosets by Hung et al. (2015). As such, we have adopted the terminology used in their manuscript: occipital (O; V2/V3), posterior ventral (PV; V4/TEO), posterior dorsal (PD; FST), middle dorsal (MD; caudal TE), and anterior dorsal (AD; rostral TE). In addition to these face-selective patches in temporal lobe, we also found clear peaks related to the directed gaze videos in anterior cingulate (at the confluence of 8b, 32, and 24) and lateral frontal cortex (45/47L). Subcortically, SC, Pul, Hipp, and MD also showed face selectivity when contrasting the directed and averted gaze conditions.

**Figure 3.**
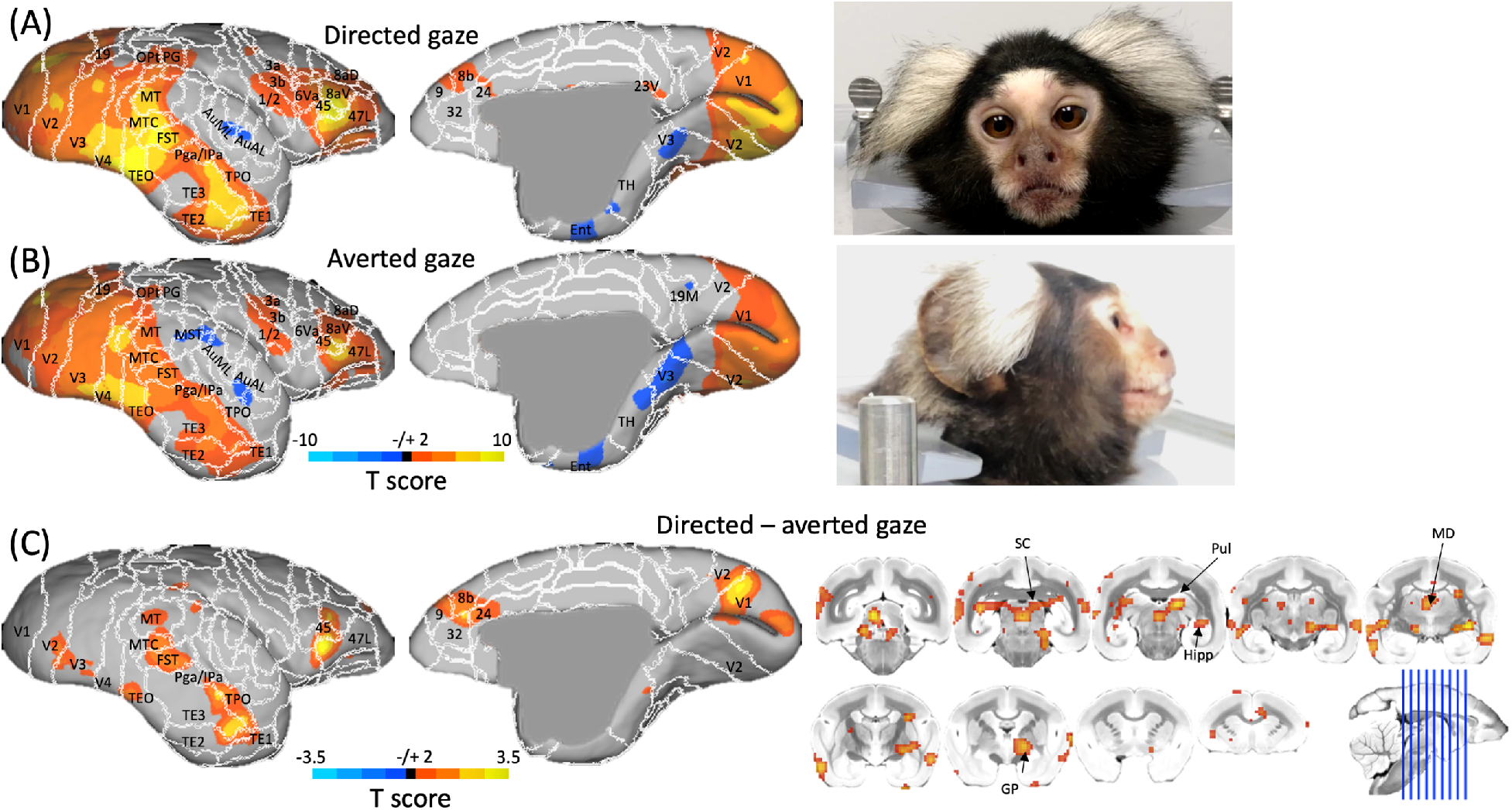
Group functional topology comparisons between directed and averted gaze video conditions displayed on a marmoset cortical surface. (A) shows group topologies for directed gaze videos and (B) shows topologies for averted gaze videos, with stills of representative stimuli to the right of each surface. (C) shows the contrast between the directed and averted gaze conditions, with volumetric display (to show subcortical activation) to the right of the medial and lateral surfaces.

### Resting-state seed analysis

With the frontal face patches being a novel finding of the task-based experiment described above (i.e., anterior cingulate and lateral prefrontal cortex differentially activating for directed gaze faces), we sought to corroborate the functional connectivity between the temporal face patches and those found in frontal cortex. As such, we calculated functional connectivity between the face patches (AD, MD, PD, PV) and the rest of the brain as shown in Figure 4A. Indeed, AD, MD, and PD all connected (with varying extents) to the 8b/24 anterior cingulate cluster and the lateral frontal cortex cluster (45/47). Generally, the anterior face patches (MD and AD) connected most strongly with anterior cingulate cortex, whereas as the more posterior face patches (PV and PD) strongly connected to lateral frontal cortex. PV connected strongly with lateral frontal cortex, but not anterior cingulate cortex.

**Figure 4.**
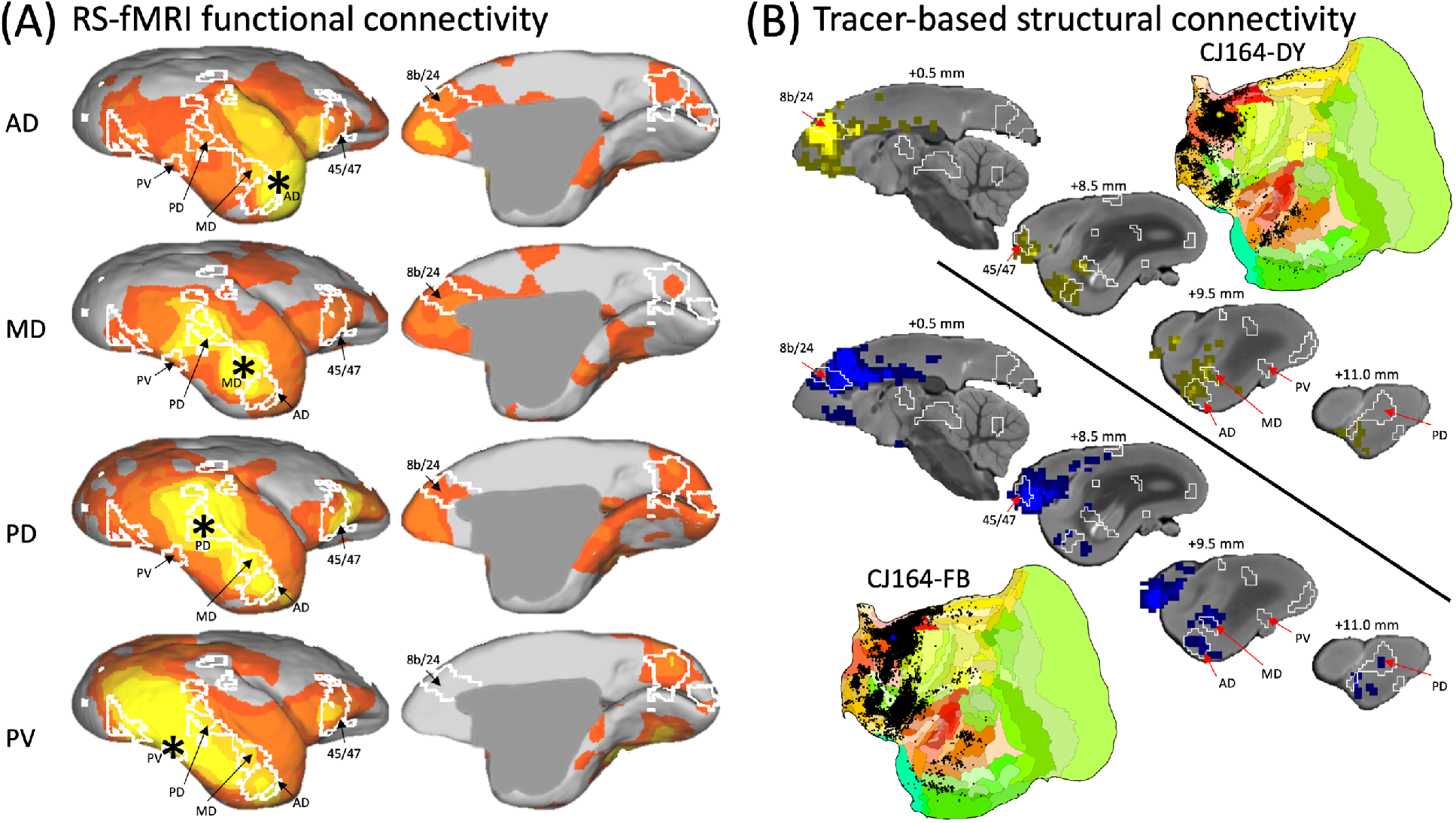
Functional and structural connectivity of marmoset face patches. (A) shows RS-fMRI based functional connectivity of four temporal face patches with the rest of the brain. (B) shows the results of two retrograde tracer injections which were proximal to our anterior cingulate cortex face patch. Surface flat map representations of these tracer injections (downloaded from marmosetbrain.org) are also shown. In both (A) and (B) white lines show the functional clusters found by comparing the directed and averted gaze conditions for reference (i.e., the topology shown in Figure 3C).

### Comparison with tracer-based cellular connectivity

With cortical tracer injections publicly available (Majka et al., 2020) we were also able to compare our findings with structural connectivity in marmosets. As shown in Figure 4B, injections proximal to the face selective anterior cingulate cortex cluster (8b/24) show clear connectivity with lateral frontal cortex (45/47). Further, consonant with our functional connectivity analysis, anterior cingulate cortex injections show strong connectivity with the anterior face patches (AD and MD), but weaker connectivity with the posterior face patches (PD and PV).

### Eye tracking

Distributions representing proportion of saccades by saccade amplitude and proportion of fixations by duration are shown in Figure 5. Separate repeated measure ANOVAs were performed for saccade amplitudes and fixation durations with the factor of conditions (4 levels: fixation, scrambled face, directed gaze, averted gaze). For saccade amplitudes, a significant effect of condition was observed, *F*(3,12) = 8.02, *p* = .00, *MSE* = .23, *η*_*p*_^*2*^ = .67, where saccade amplitudes were significantly longer in the averted gaze condition than in the fixation or scrambled face conditions. However, no significant differences between conditions were observed after correcting for multiple comparisons. Similarly for fixation durations, a significant effect of condition was observed, *F*(3,12) = 4.98, *p* = .02, *MSE* = 86.28, *η*_*p*_^*2*^ = .55, where fixation duration was longer for the fixation condition than for the averted gaze condition but pairwise comparisons revealed no significant differences between conditions after corrections. Overall, these results suggest that the cortical topologies were likely not directly driven by differences in saccade number or amplitudes.

**Figure 5.**
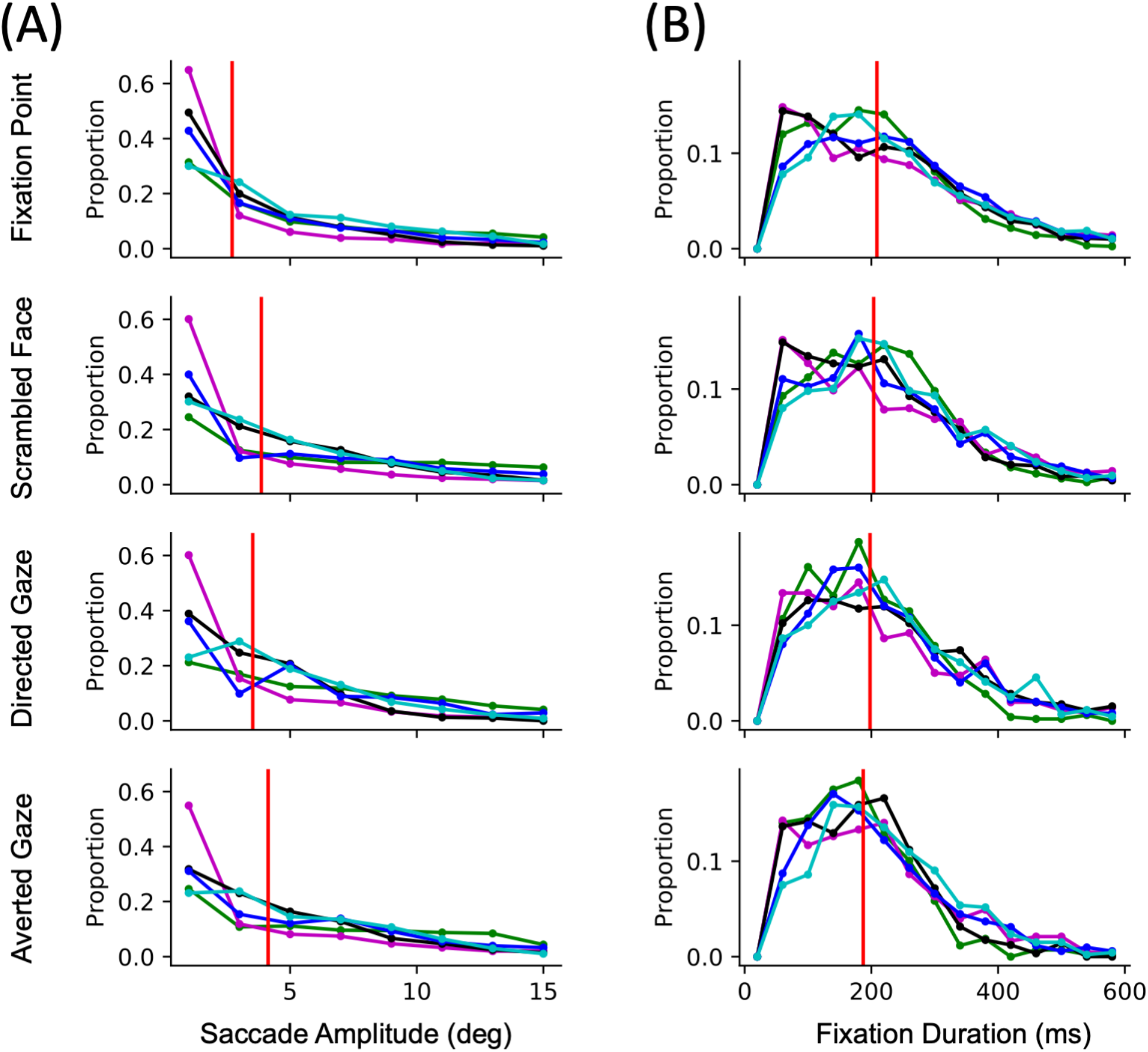
Distributions of saccades by saccade amplitude in visual degrees (A) and fixations by fixation duration in ms (B) for each condition separately for 5 marmoset subjects. Red lines represent the group median value.

## Discussion

In this study, we were interested in determining whether New World marmosets show face-selective patches in frontal cortex, as has been demonstrated in Old World macaques (Tsao et al., 2008a, 2008b; Shepherd and Freiwald, 2018). To do so, we presented videos of marmoset faces during fMRI acquisition at ultra-high field. Similar to macaques, marmosets showed differential activation in anterior cingulate cortex and lateral frontal cortex when viewing videos of conspecific faces with directed gaze (i.e., direct eye contact) when compared to averted gaze videos (Shepherd and Freiwald, 2018). To corroborate the connectivity of these frontal face patches, we also compared our task-based fMRI results with RS-fMRI based functional connectivity and tracer-based structural connectivity. Both analyses demonstrated strong connectivity between the temporal face patches and the frontal face patches, with the anterior and posterior patches differentially connected to medial (8b/24) and lateral frontal cortex (45/47), respectively. Overall, these findings suggest that marmosets do indeed possess a face processing circuitry that extends into frontal cortex and likely supports socially relevant processing of faces.

When comparing patterns of activation elicited from videos of marmoset faces, our results are remarkably similar to those shown in a previous marmoset fMRI study (Hung et al., 2015) which presented photos of marmoset faces, bodies, objects, or scrambled versions of these photos and found face selectivity in six occipitotemporal patches (O, PD, PV, MD, AD, and MV). We corroborate their results by eliciting all of these patches with marmoset face videos, with the exception of MV – in fact, this group also did not see this patch with fMRI acquisition, but rather used electrocorticography arrays to index activity in this ventrolateral region. Here, we likely also did not see MV in our topologies because of the low signal-to-noise ratio in this area (see Schaeffer et al., 2019a for receive coil design and signal-to-noise topologies).

The temporal face selective patches in marmosets shown here are comparable to those found in humans and macaques (as reviewed in Weiner and Grill-Spector, 2015). In humans, these regions exist along the ventral temporal surface, which is a slightly different organization than in macaques, which show patches that are more dorsally located, around the superior temporal sulcus (Tsao et al., 2008a). These Old World primate species, however, also show face selective patches in frontal cortex – to date, this had yet to be demonstrated in New World marmosets. Here, as shown in Figure 3, we demonstrate that marmosets do indeed show face selectivity in anterior cingulate and lateral frontal cortex. These patterns were obtained by contrasting patterns of activation elicited from videos of marmosets with directed gaze (Figure 3A) versus marmosets looking toward the left or right (i.e., averted gaze; Figure 3B). As previously demonstrated in macaques (using a similar conspecific video set), the patch in anterior cingulate cortex (8b/24) was specific to the directed gaze condition, suggesting that this patch is involved in socially relevant face processing.

The lateral prefrontal patch (peak between 45 and 47) shown here, however, was elicited in both directed and averted conditions, albeit to a stronger degree in the directed gaze condition. We hypothesize that the dorsal part of this functional cluster, especially that reaching into area 8aV, is related to small eye movements (i.e., making saccades to salient features of the videos), rather than being face-specific *per se.* The lateral portion of this cluster, however, is likely face specific – we have recently demonstrated topologies related to saccadic eye movements in marmosets using both microsimulation (Selvanayagam et al., 2019) and task-based fMRI (Schaeffer et al., 2019b) – when directly overlaying these patterns, it seems that the peak in area 45/47 shown in Figure 3C is likely too far ventrolateral to be related to eye movements. Further, neither the saccade amplitude nor fixation duration differed between these conditions (Figure 5). Therefore, this patch in lateral frontal cortex is likely face selective. Given that this patch is present in both macaques and humans, we hypothesize that this patch could be the marmoset homolog to these regions (Tsao et al., 2008a, 2008b; Schwiedrzik et al., 2015; Shepherd and Freiwald, 2018).

Evidence for face selectivity in anterior cingulate cortex and lateral frontal cortex patches was further substantiated by both functional and structural connectivity between these regions and the temporal face patches. To index functional connectivity, we utilized our ultra-stable (~150 μm maximum head motion), fully awake RS-fMRI dataset acquired at 9.4 Tesla. We seeded four of the face patches (AD, MD, PD and PV, but did not seed O due to concerns with low signal, see Schaeffer et al., 2019a on this issue) and found strong connectivity between AD, MD, and PD with the anterior cingulate cortex face patch (Figure 4A). Interestingly, the anterior face patches (MD and AD) connected most strongly with anterior cingulate cortex, whereas the more posterior face patches (PV and PD) strongly connected to lateral frontal cortex. This pattern was also present when we examined tracer-based structural connectivity (Figure 4B), with two separate retrograde tracer injections proximal to the anterior cingulate face patch showing clear connectivity with the AD and MD patches. Further, these anterior cingulate cortex injections also showed strong connectivity with the lateral frontal face selective patch elicited via the directed gaze videos. Taken together, these results suggest that the anterior cingulate cortex and lateral frontal cortex face patches are part of a cortical face processing network in the marmoset brain.

Our results also provide evidence for the subcortical constituents of a face processing network in marmosets. With our custom cortex-oriented receive coil (Schaeffer et al., 2019a) we were not equipped to be maximally sensitive to subcortical regions, but the general pattern found here is quite similar to that shown in Old World primates (Johnson, 2005; Mosher et al., 2014; Kuraoka et al., 2015; Schwiedrzik et al., 2015; Sliwa and Freiwald, 2017). When contrasting the social and non-social scrambled conditions, we found differential activation in SC, Hipp, Pul, MD, and Amy. Face selectivity in regions such as SC, Pul, Hipp and Amy is intriguing and our findings certainly warrant further investigation on this subject, perhaps with more sensitive electrophysiological techniques that have been used to show subcortical face selectivity in macaques (Mosher et al., 2014; Nguyen et al., 2014). The fMRI results here may serve as a starting block for these more invasive explorations of face selective patches across the marmoset brain.

In summary, we report evidence from task-based fMRI, RS-fMRI, and tracer based structural connectivity for a face processing network in New World marmosets which includes at least two face selective patches in frontal cortex. We demonstrated that, as in Old World primates, these frontal face patches are differentially sensitive to social interaction (i.e., direct eye contact), suggesting a high-level, top-down control social processing network in the marmoset brain. Therefore, our results suggest that the origin of this frontal network specialized for social face processing predates the separation between Old and New World primates around 35 million years ago (Schrago and Russo, 2003). These results give further credence to the marmoset as a viable preclinical modelling species for studying human social disorders.

## Author Notes

Support was provided by the Canadian Institutes of Health Research (FRN 148365, FRN 353372), a Brain Canada Platform Support Grant and the Canada First Research Excellence Fund to BrainsCAN. We wish to thank Cheryl Vander Tuin, Whitney Froese, Kathrine Faubert, and Miranda Bellyou for surgical assistance, animal preparation and care and Dr. Alex Li for scanning assistance.

